# KPNA2 promotes cellular proliferation and inhibits apoptosis in the Saos-2 osteosarcoma cell line

**DOI:** 10.1101/436428

**Authors:** Dinglu Wei, Qiaofeng Ge, Xiaojuan Sun, Long Zhang, Jia Li, Chenglong Chen, Zhi Lv

## Abstract

Karyopherin α2 plays a critical role in tumorigenesis and tumor progression. However, nothing is currently known about the effects of KPNA2 on osteosarcomas. This study aimed to investigate differential KPNA2 protein and mRNA expression in human osteosarcoma tumor cells and normal bone tissue. We also sought to determine whether KPNA2 can influence the proliferation and apoptosis of the Saos-2 osteosarcoma cell line. Immunohistochemistry (IHC) was used to investigate KPNA2 protein expression. Real-time quantitative PCR (qPCR) was used to detect levels of KPNA2 mRNA expression, and lentivirus-mediated short-hairpin RNAs (shRNAs) were used to knock down KPNA2 expression in Saos-2 cells. The MTT assay and multiparametric high-content screening (HCS) were used to measure cell proliferation and growth, respectively. Flow cytometry was conducted to detect cell cycle distribution and apoptosis. The results revealed significantly higher KPNA2 expression levels in osteosarcoma tissues than in normal bone tissues; furthermore, KPNA2 mRNA was also highly expressed in three osteosarcoma cell lines. After transducing Saos-2 cells with KPNA2-shRNA lentivirus, the proliferative rate was notably decreased compared to that of the negative control (NC) lentivirus group (P<0.05). Flow cytometry results indicated that KPNA2 may arrest cell cycle progression and regulate the growth of these cells. The results for apoptosis indicated an apoptotic rate of 13.38±0.48% in KPNA2-shRNA cells, which was significantly higher than the rate for cells in the control group (5.13 ±0.33%). Therefore, this study showed that KPNA2 is highly expressed in osteosarcoma tissues and that reduced KPNA2mRNA levels inhibited proliferation and promoted apoptosis in an osteosarcoma cell line.

## Introduction

Osteosarcoma is the most common primary bone malignancy and often manifests in the long bones, i.e., the distal femur and proximal tibia (1, 2). Osteosarcoma is the leading cause of cancer-related death in childhood and adolescence with an incidence of 5/1,000,000 per year in the United States (2). Currently, surgical resection and chemotherapy are the primary treatments for osteosarcoma; however, patients suffering from chemoresistance or metastatic disease still have poor outcomes (3–5). Thus, developing new treatments aimed at specific molecular targets and acquiring deep exploration of the molecular mechanisms of the currently available therapeutic strategies are urgently needed to combat the tumorigenesis of osteosarcomas.

The dysfunction of nuclear transport machinery has been described in several tumor types and plays an important role in tumorigenesis and tumor progression (6). Cargo proteins must pass through large nuclear pore complexes (NPCs) within the nuclear membrane to complete nucleocytoplasmic transport (7). The transport of macromolecules larger than 40 kDa cannot cross the nuclear membrane without the mediation of karyopherins—key carrier proteins (7–9). The karyopherin family comprises import and export factors that participates part in several nuclear transport pathways into and out of the nucleus (9). Karyopherin α2 (KPNA2) is a member of the karyopherin α family, which also includes importin α-1 and RAG cohort 1 (8). KPNA2 has a mass of approximately 58 kDa and contains 529 amino acids (7) that compose three domains: an N-terminal hydrophilic importin β-binding domain, a central hydrophobic region consisting of 10 armadillo (ARM) repeats (which bind the cargo’s NLS) and a short acidic C-terminus with no reported function (7).

In recent years, aberrant expression of KPNA2 have been reported in a variety of cancers, including bladder cancer (10), lung cancer (11), liver cancer (12), ovarian cancer (13), melanoma (14), esophageal cancer (15), prostate cancer (16) and colon cancer(21). Some studies revealed KPNA2 as a potential biomarker for multiple types of malignancies, and Dahl et al. reported the prognostic value of elevated KPNA2 expression in breast cancer (17). However, there are currently no reports about the expression and the role of KPNA2 in osteosarcoma.

In this study, we first analyzed expression of KPNA2 in osteosarcoma tumor samples using immunohistochemistry (IHC) and evaluated the mRNA levels of KPNA2 in osteosarcoma cell lines using real-time quantitative PCR (qPCR). Then, lentivirus-mediated transduction of shRNA was performed to knock down KPNA2 mRNA expression in the Saos-2 osteosarcoma cell line. Western blot analysis was used to detect the protein levels of KPNA2 and PCNA in infected Saos-2 cells. Finally, we evaluated the effect of silencing KPNA2 on *in vitro* cellular characteristics such as cell cycle, apoptosis, and proliferation.

## Materials and methods

### Clinical specimens

Thirty five osteosarcoma samples and five normal bone tissues were obtained from the Department of Pathology of Shanxi Medical University Second Affiliated Hospital from 2005 to 2015. The patients from whom the samples were obtained had not received any treatment before tumor resection. This research was authorized by The Ethics Committee of Shanxi Medical University.

### Immunohistochemistry

A 4% formaldehyde solution was used to fix all tissue samples. All samples were then embedded in paraffin blocks and cut into 5-μm sections. All sections were placed on glass slides, dewaxed with xylene and rehydrated with alcohol, followed by heat-induced antigen retrieval using 10 mmol/L citrate buffer. Methanol containing 3% hydrogen peroxide was used to inhibit endogenous peroxidase activity. The slides were blocked with normal serum for 30 min and then treated with an anti-KPNA2 antibody (1:100) (Boster Biotechnology Company, Wuhan, China) overnight at 4°C. Phosphate-buffered saline (PBS) replaced the primary antibody as a negative control. Each slide was washed three times and then treated with the secondary antibody (anti-mouse, Boster Biotechnology Company, Wuhan, China) for 15 min and stained with diaminobenzidine.

The percentage of stained cells and intensity scores were used to obtain a total score for each slide. The scores ranged from 0 to 7, with 0-4 indicating negative expression and 4-7 indicating positive expression.

### Cell culture

The human osteosarcoma cell lines Saos-2, HOS, and MG63 and human chondrocyte cell C-28 were purchased from CBTCCCAS (Shanghai, China) and cultured in Roswell Park Memorial Institute (RPMI) 1640 medium supplemented with 2 mM L-glutamine, 100 μg/ml streptomycin, 100 U/ml penicillin and 10% fetal bovine serum (Gibco, Grand Island, NY, USA). All cells were cultured in a humidified atmosphere of 5% CO2 at 37°C

### Real-time polymerase chain reaction (RT-PCR)

Total RNA from three osteosarcoma cell lines, C28 cells and Saos-2 cells after infected with the KPNA2-shRNA lentivirus were extracted by using TRIzol reagent (Invitrogen, USA). Then, cDNA was synthesized from 2 μg of total RNA per sample using a Reverse Transcription System according to the manufacturer’s instructions and was used as the template to quantify the relative mRNA content. The qPCR cycling conditions were as follows: an initial denaturation at 95°C for 15 s followed by 45 cycles at 95°C for 5 s and 60°C for 30 s. The sequences of the primer pairs used in the PCR were as follows: GAPDH, forward 5’- TGACTTCAACAGCGACACCCA-3’, reverse 5-CACCCTGTTGCTGTAGCCAAA-3’; and KPNA2, forward 5’- TGTGGTAGATGGAGGTGC-3’, reverse 5’- GAGCCAACAGTGGGTCA-3’. The gene expression data were analyzed by using the 2-ΔΔCt method.

### *Lentivirus vectors for KPNA2 short-hairpin RNA* and infect the human osteosarcoma *Saos-2 cell line*

We used the PSCSI-GFP lentiviral plasmid to express short-hairpin RNAs (shRNAs) targeting the KPNA2 ORF sequence (GenBank no. NM_002266). The sequences 5’- ACCTCTGAAGGCTACACTT-3’ was cloned into the PSCSI-GFP plasmid prior to generating the lentivirus (GeneChem, Shanghai, China) and a non-specific targeting sequence (GeneChem, Shanghai, China) and was cloned into the PSCSI-GFP plasmid prior as a negative control (NC). Human renal epithelial 293T cells were used for packaging the viral particles for obtaining the KPNA2-shRNA lentivirus and a negative control (NC) lentivirus. Western blotting was used to detect the interference efficiency. GFP expression was observed by fluorescence microscopy 3 days after infection, the mRNA and protein expression levels of KPNA2 were measured by qPCR and Western blot analysis to determine the knockdown efficiency 5 days after infection.

### Western blot analysis

Saos-2 cells were infected with either KPNA2-shRNA lentivirus or NC lentivirus, and total cellular protein was harvested using RIPA lysis buffer (Pierce, Shanghai, China) at 36 h after infection. Total proteins were separated by 10% sodium dodecyl sulfate-polyacrylamide gel electrophoresis (SDS-PAGE) and transferred to polyvinylidene difluoride (PVDF) membranes. The PVDF membranes were incubated with mouse anti-KPNA2 (1:2000, Sigma, USA), mouse anti-PNCA (1:2000, Sigma, USA) and mouse anti-ß-Actin (1:2000, Santa Cruz Biotechnology, USA) primary antibodies at 4°C overnight. Goat anti-mouse IgG (1:2000, Santa Cruz Biotechnology, USA) was used as the secondary antibody, and immuno-complexes were detected using Pierce™ ECL Western Blotting Substrate.

### Cell growth assay

Multiparametric high-content screening (HCS) was used to measure cell growth. Briefly, after infection with either NC lentivirus or KPNA2-shRNA lentivirus, Saos-2 cells were cultured for 72 h and harvested at the logarithmic growth phase. The resulting cell suspensions were seeded in 96-well plates at 2,000 cells per well and incubated at 37°C in a humidified atmosphere containing 5% CO_2_ for 5 days. Cell growth in each plate was measured using ArrayScan™ HCS software (Cellomics Inc.). Cell images were captured using ArrayScan™ HCS software once a day for 5 days. The number of cells per well was quantified using ArrayScan™ HCS software. The cell numbers in each well were used to generate cell growth curves for each condition.

### MTT cell viability assay

Briefly, Saos-2 cells were infected with either the NC lentivirus or KPNA2-siRNA lentivirus and then collected during the logarithmic growth phase. The resulting cell suspensions were seeded at 2000 cells per well in 96-well plates. Cells were incubated at 37°C for 1, 2, 3, 4, and 5 days. At the indicated time points, cells were washed twice with PBS and treated with 20 μL of MTT solution (5 mg/mL) per well. After a 4-hour incubation, the supernatants were removed, and 100 μL of dimethyl sulfoxide (DMSO) was added to each well to solubilize the formazan salt. The optical density (OD) was measured at 490 nm by using a microplate reader (Tecan infinite, Austria). Each experiment was performed in triplicate and repeated three times.

### Flow cytometry

Flow cytometry was conducted to detect cell cycle distribution and Cell apoptosis. After infection with either KPNA2-shRNA lentivirus or negative control lentivirus, cells were cultured in 6-well plates. Upon reaching 80% confluence, the cells were collected by trypsinization, washed once with PBS, and fixed in 70% alcohol for 2 h. For the cell cycle analysis, the fixed cells were washed with D-Hanks and stained with propidium iodide (PI) buffer containing 40× PI stock (2 mg/ml), 100× RNase stock (10 mg/ml) and 1× D-Hanks stock at a dilution ratio of 25:10:1,000. At least 1 × 10^4^ cells of each sample were analyzed by flow cytometry (Guava easyCyte HT, Millipore, USA) to determine the cell cycle phase distribution. For the cell apoptosis analysis, the cells were resuspended in 200 μL of 1× staining buffer containing 10 μL of annexin V-APC followed by an incubation at room temperature in the dark for 10–15 min. Afterwards, the cells were analyzed by using flow cytometry (Guava easyCyte HT, Millipore, USA). All experiments were performed in triplicate.

### Statistical analysis

Student’s t-test and the Kruskal-Wallis test were used for raw data analysis. Statistical analysis was performed using the SPSS software package. All data are expressed as the mean ± standard error of the observations. P < 0.05 was considered to be the significance level. All experiments were repeated in triplicate.

## Results

### Expression of KPNA2 in osteosarcoma

IHC was used to investigate KPNA2 protein expression in 35 human osteosarcoma tumor tissues and 5 normal bone tissue samples. RT-PCR was used to measure KPNA2 mRNA levels in Saos-2, MG63, HOS and C-28 cells. As shown in Figure 1A:, among the 35 cancerous tissue samples, positive KPNA2 immunostaining was observed in the nuclei of the osteosarcoma tumors; 30 samples (88.6%) showed positive staining for KPNA2, but 5 (11.4%) showed negative staining, while all the 5 normal bone tissue sample were negative. As shown in Figure 1B, compared with the human chondrocyte cell line C-28, all osteosarcoma cell lines expressed higher levels KPNA2 mRNA, with the highest levels observed in Saos-2 cells. Therefore, we chose Saos-2 cells for all subsequent experiments.

**Figure 1A.**
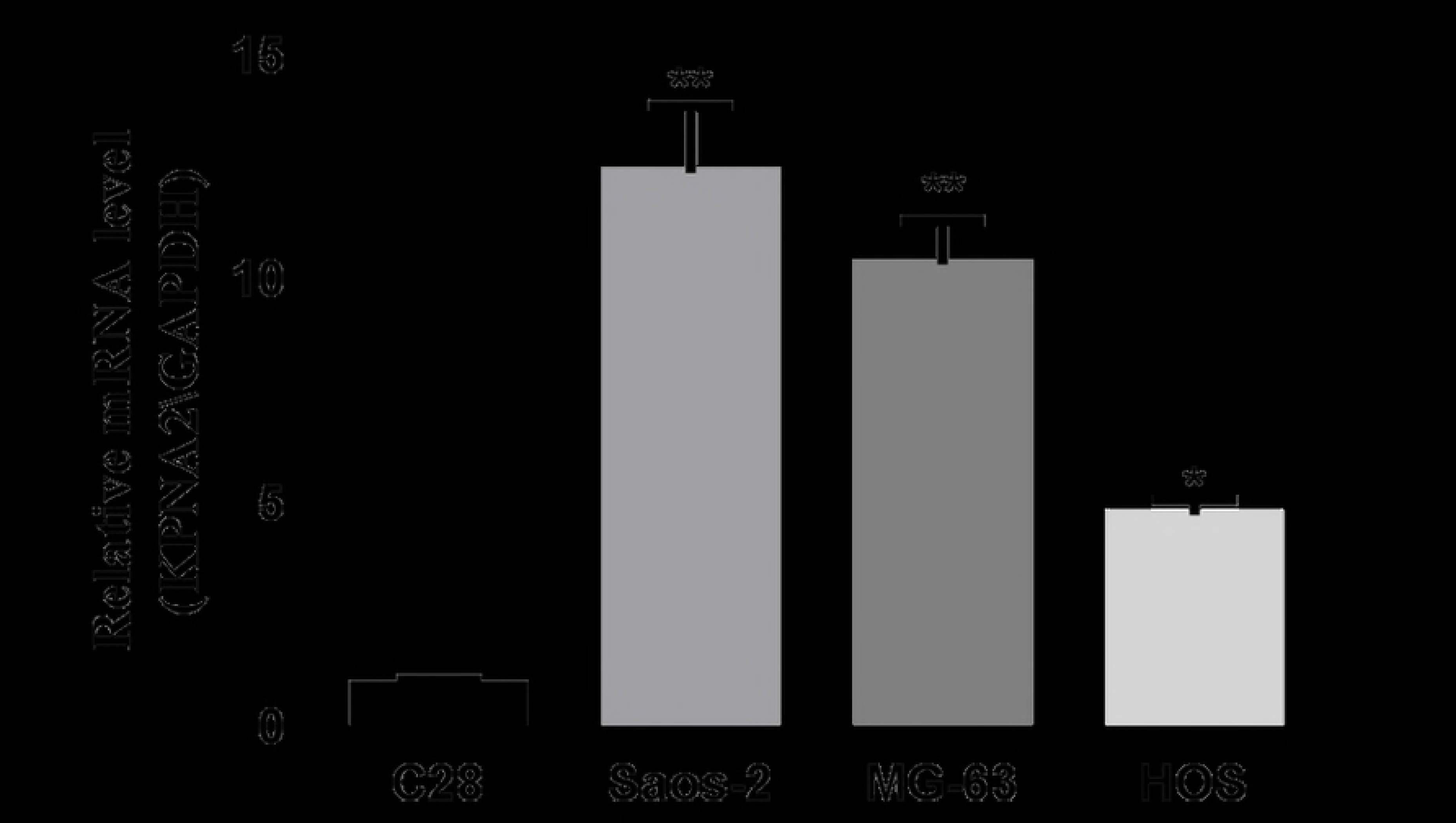
Immunohistochemical staining of KPNA2 in osteosarcoma and normal bone tissues. The brown grains indicate a positive signal. The brown grains were primarily localized in the nucleus of tumor cells. Fig. 1A-a, Negative KPNA2 expression in normal tissue. Fig. 1A-b, Negative control to demonstrate the specificity of the antibody. Fig. 1A-c, Positive KPNA2 expression in osteosarcoma tissue. Original magnification, 200× in a-c. The scale bar represents 100 μm.

**Figure 1B.**
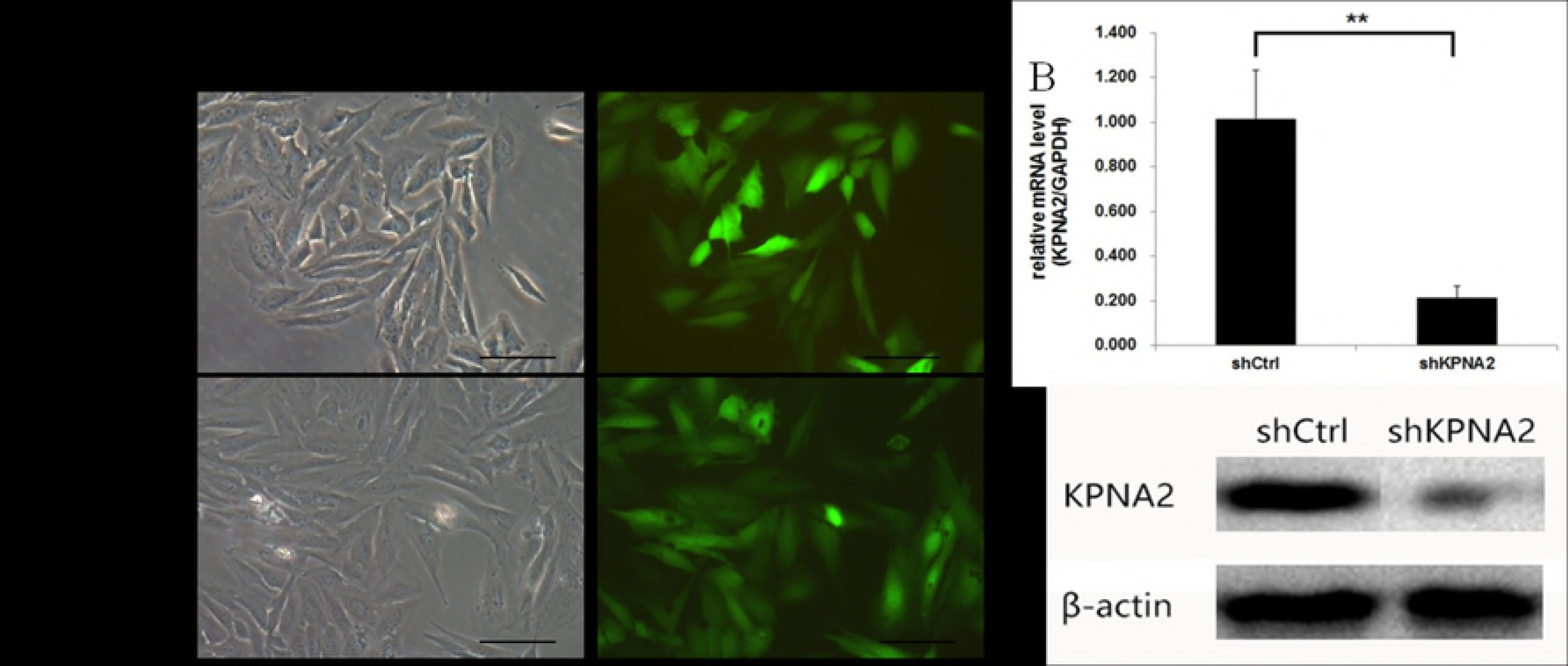
KPNA2 mRNA expression levels were detected by RT-PCR in three osteosarcoma cell lines, GAPDH served as an internal control.**P < 0.01 when Saos-2 or MG-63cells compared to C28cells. *P < 0.05 when HOScells compared to C28 cells.

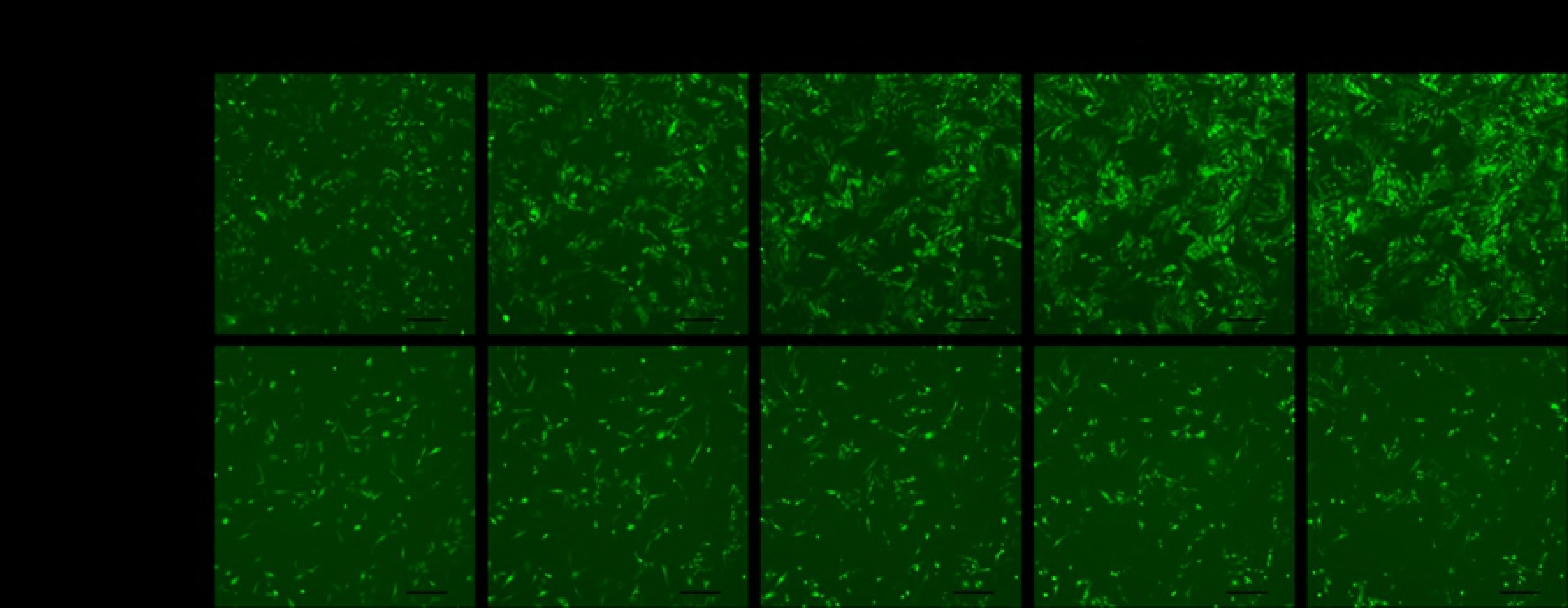

These data indicate that the KPNA2 protein and mRNA are overexpressed in osteosarcoma.

### KPNA2 expression was efficiently inhibited by KPNA2-shRNA lentivirus transduction in Saos-2 cells

To investigate the role of KPNA2 in osteosarcoma, we first needed to knock down KPNA2 levels in the human osteosarcoma cell line Saos-2. As described in the methods section, either shRNA specifically targeting the KPNA2 ORF sequence or a non-specific sequence was cloned into the PSCSI-GFP plasmid to generate lentivirus. Saos-2 cells were then infected by either KPNA2-shRNA lentivirus or NC lentivirus (both of which express GFP), and GFP expression in the cells was observed by fluorescence microscopy.

As shown in Figure 2A, more than 80% of the cells expressed GFP at 3 days after infection, indicating that the infection efficiency of the lentivirus was greater than 80%. At 5 days after infection, the KPNA2 mRNA levels in Saos-2 cells were detected by qPCR. As shown in Figure 2B, compared with the NC group (shCtrl), the KPNA2-shRNA group (shKPNA2) showed an obvious decrease in the KPNA2 mRNA levels, with an approximate 78% reduction. Western blotting analysis was used to detect the protein levels in the Saos-2 cells at 4 days after infection. As shown in Figure 2C, the KPNA2 protein levels were greatly reduced in cells expressing KPNA2-shRNA compared with those in the negative control group, indicating that KPNA2-shRNAs effectively knocked down KPNA2 protein expression.

**Figure 2.**
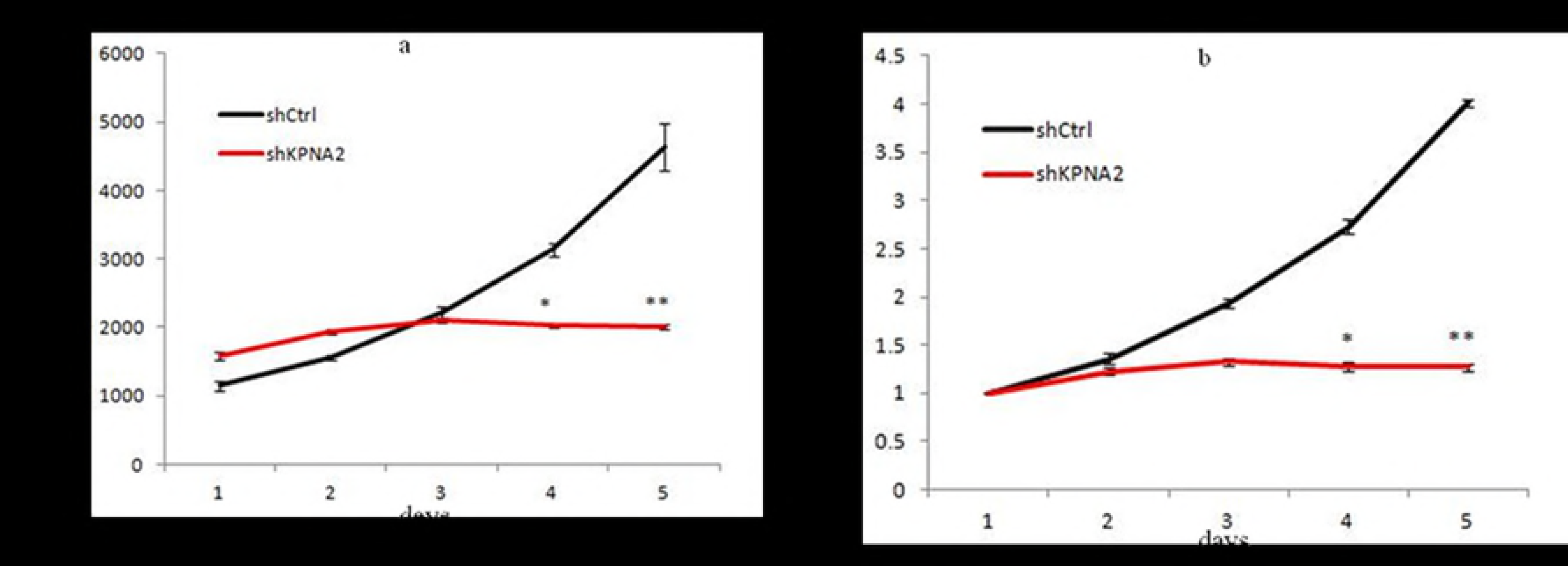
Representative images of Saos-2 cells infected with KPNA2-siRNA or NC siRNA (magnification ×200, scale bars 100 μm) Figure 2-A. The green GFP fluorescence reflects the efficiency of lentiviral infection. Figure 2-B KPNA2 mRNA levels in Saos-2 cells were detected by qPCR 5 days after infection. GAPDH was used as an internal control. The data are presented as the mean ± SD, **P < 0.01

### The effect of KPNA2 effect on cell proliferation in human osteosarcoma cells

To explore the function of KPNA2 in regulating melanoma cell proliferation, Saos-2 cells expressing either KPNA2-shRNA or NC siRNA were seeded in 96-well plates, and the number of cells was counted each day for 5 days using Cellomics (Figure 3A). The cell growth rate was assessed using the following equation: cell count on the Nth day/cell count on the 1st day, where N is 2, 3, 4, or 5. The results showed that downregulation of KPNA2 decreased the total number of cells (Figure 3B-a) and slowed the growth rate (Figure 3B-b). The relationship between KPNA-2 and the proliferative ability of the cells was also determined using the MTT assay. As shown in Figure 3C, the proliferative rate of the KPNA2-shRNA group was markedly decreased when compared with that of the NC shRNA (shCtrl) group. The protein level of PCNA can also reflect the cell proliferation ability; therefore, Western blotting was used to detect the PCNA protein levels in Saos-2 cells 4 days after infection. As shown in Figure 3D, the PCNA protein levels were greatly reduced in the cells expressing KPNA2-shRNA compared with those in the negative control group, indicating that the cell proliferations ability of the KPNA2-shRNA group was effectively decreased.

**Figure 3.**
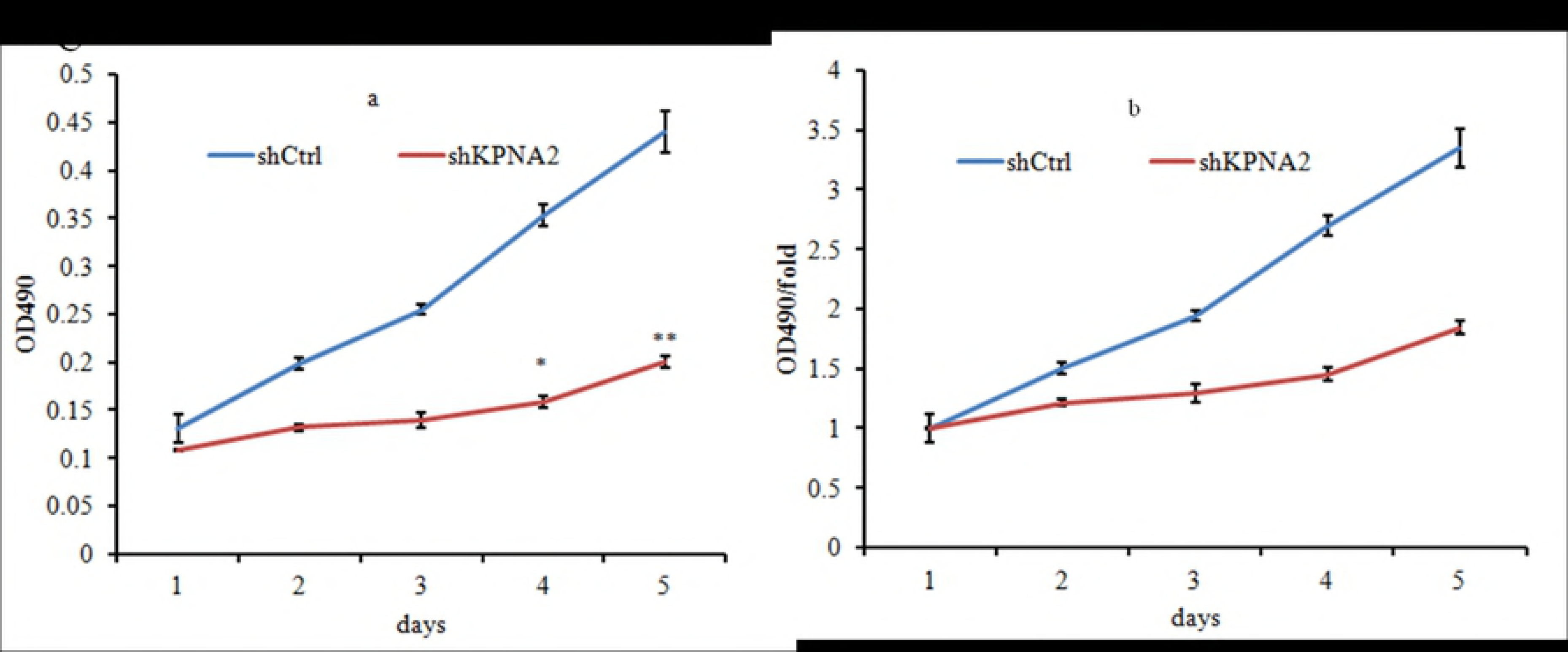
Knockdown of KPNA2 inhibited cell proliferation. Figure3A Representative raw images of Saos-2 cell growth at different time points after lentivirus infection. Figure3B-a The number of cells was quantified daily for 5 days. (NC vs KPNA2-shRNA, **P < 0.01, *P < 0.05). Figure3B-b, the growth rate was nitored on the 2nd, 3rd, 4th and 5th days using an assay (NC vs KPNA2-shRNA, **P < 0.01,*P < 0.05). Figure3C-a, the optical density (OD) was measured using a microplate reader at 490 nm. Figure3C-b, the proliferation rate of the KPNA2-shRNA group was notably reduced, compared with that of the control group (NC vs KPNA2-shRNA, **P < 0.01,*P < 0.05).

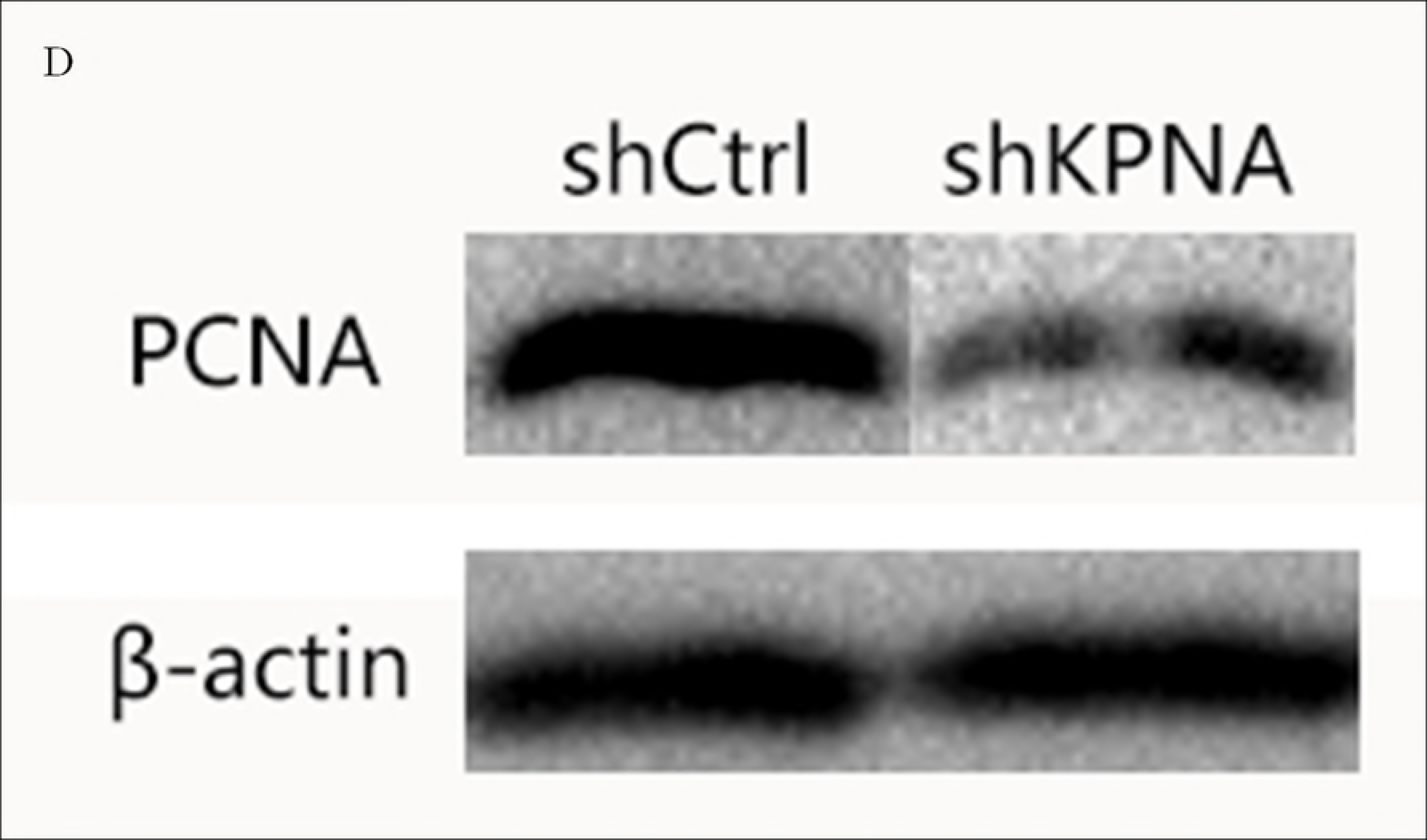

### Knockdown of KPNA2 arrested the cell cycle progression

As is commonly known, cancer cells exhibit characteristic dysregulation of the cell cycle. Therefore, we wanted to explore the effect of KPNA2 on cell cycle progression in the Saos-2 human osteosarcoma cell line. The cell cycle distribution of Saos-2 cells after KPNA2 knockdown was analyzed by flow cytometry (Figure 4A). As shown in Figure 4B, the control group showed 53.50% in G0/G1 phase, 38.49% in S phase, and 8.01% in G2/M phase; however, the distribution of cells in the KPNA2-shRNA group was 35.86%, 53.77%, and 10.36%, respectively. Compared with that of the shCtrl group, the KPNA2-shRNA group displayed a remarkable decrease in G0/G1 phase (35.86% vs. 53.50%) and a significant increase in S phase (38.49% vs 53.77%). These results suggest that KPNA2 may arrest Saos-2 cell cycle progression and regulate Saos-2 cell growth.

**Figure 4A.**
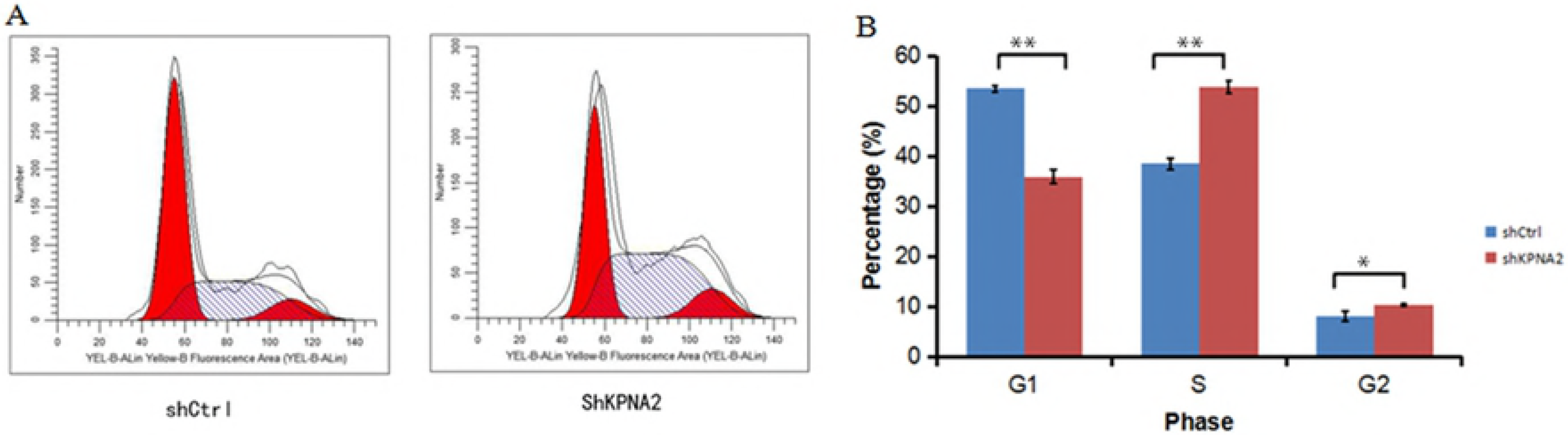
The effect of KPNA2 on cell cycle progression was determined by flow cytometry.

**Figure 4B.**
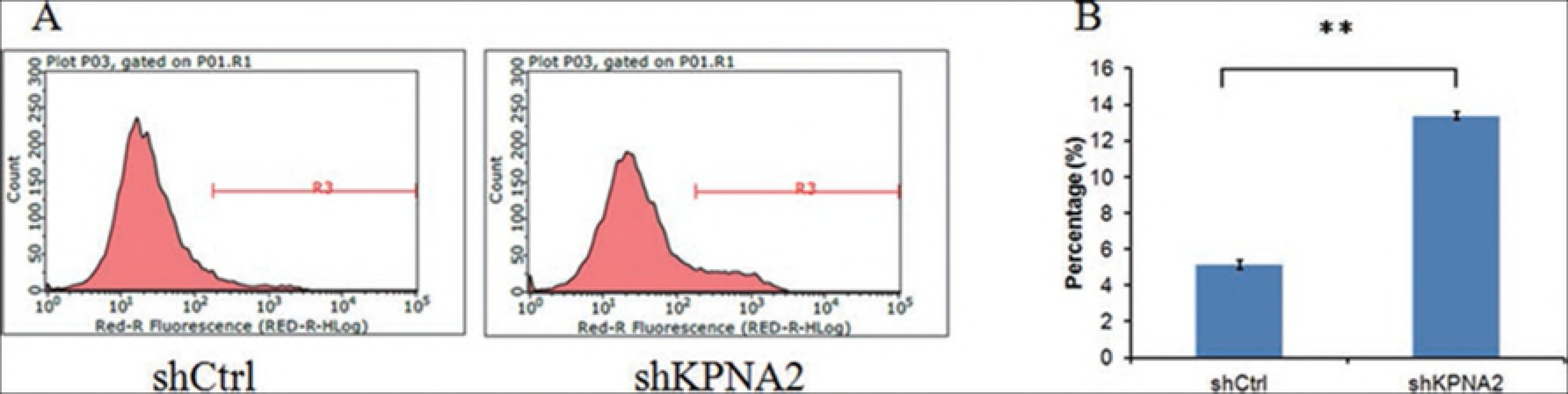
The results showed that compared with the control group, the KPNA2-shRNA group displayed a remarkable increase in the percentage of cells in the S phase (**P < 0.01,*P < 0.05).

### Knockdown of KPNA2 induced cell apoptosis in human osteosarcoma cells

Given the findings above, we sought to determine whether KPNA2 affects Saos-2 cell apoptosis. At 5 days after infection, the apoptosis rate of Saos-2 cells was determined by staining with labeled annexin V and detecting the fluorescence with flow cytometry (Figure 5A). As shown in Figure 5B, we observed a significantly higher apoptosis rate for Sao-2 cells of the KPNA2 knockdown group than for cells of the shCtrl group (13.38 ± 0.48% vs. 5.13 ± 0.33%, P < 0.01). Therefore, this result indicated that knockdown of KPNA2 in Saos-2 cells could induce cell apoptosis.

**Figure 5.**
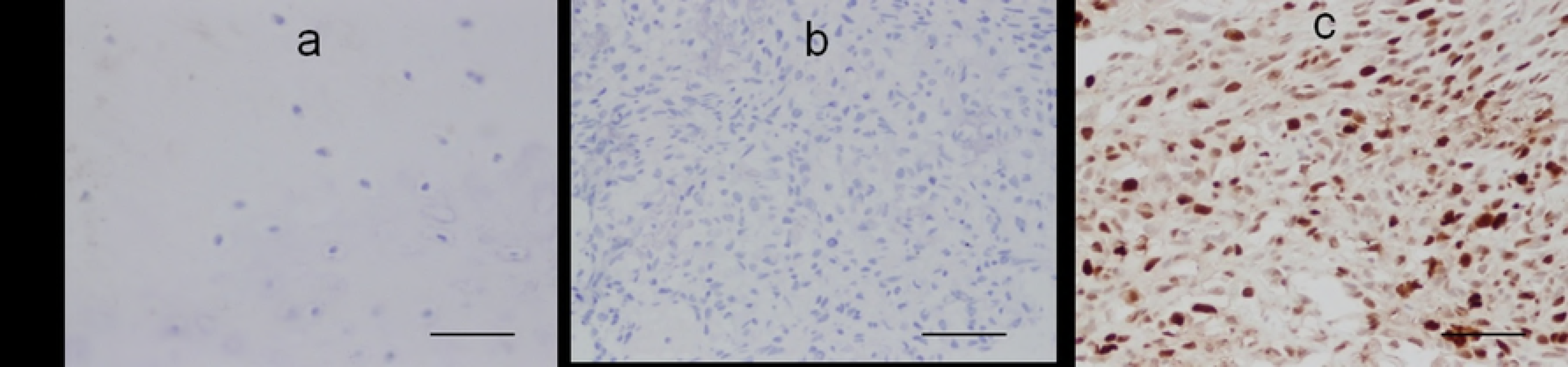
Knockdown of KPNA2 induces cell apoptosis. Figure 5A Cell death was determined by flow cytometry and Annexin V staining. Figure 5B The apoptosis rate of the KPNA2-shRNA group was 13.38 ± 0.48%,which was higher that of the control group (5.13 ± 0.33%; **P < 0.01).

## Discussion

Nucleocytoplasmic transport plays an important role in the cellular processes, which regulates gene expression, cell cycle progression, and signal transduction(18) As is commonly known, Cellular transport dysfunction is commonly observed in cancer, and tumorigenesis is often associated with increased nuclear activity in tumor cells (7). KPNA2 functions as an adaptor in the progression of cellular transport (19). In this study, our IHC staining results indicated that the level of the KPNA2 protein was significantly higher in osteosarcoma tissues compared with normal bone tissues. We explored the mRNA expression levels of KPNA2 in three osteosarcoma cell lines (Saos-2, MG63, and HOS) by qRT-PCR, and found all the three cell lines expressed high level of KPNA2 mRNA. The results further indicated, and observed that both osteosarcoma tissues and cell lines overexpressed increased levels of KPNA2. The phenomenon of KPNA2 overexpression have been observed in many malignancies (7). In the study of colon tumor Meng Zhang found that the mRNA and protein expression levels of KPNA2 were upregulated in colon tumor samples and correlated with poor survival in a large cohort of colon cancer patients (21). The similar results were also been found in endometrial cancers and gastric adenocarcinoma (22)researches. All of these studies revealed that KPNA2 may be important for tumor progression and that elevated KPNA2 expression could serve as an independent biomarker for poor survival [23].

To test the role of KPNA2 related to the cell apoptosis and cell proliferation in the osteosarcoma cell lines, we knocked down KPNA2 expression in the Saos-2 cell line by using lentiviral-mediated transduction of shRNA. Compared to control cells, the cells treated with KPNA2-shRNA showed the enhanced apoptosis rates and the reduced proliferation rates. Our results are in line with other studies in different tumors. H-Y Wang found remarkably reduced cell viability when KPNA2 was downregulated in EFO-21 and SK-OV3 cell lines. Another urothelial carcinoma study reported that KPNA2 knockdown activated the apoptosis pathway and decreased the proliferation of urothelial carcinoma cells(23). In the present study, we found that KPNA2 knockdown arrested the cell cycle progression at S phases. However, H-Y Wang reported that downregulation of KPNA2 arrested EOC cells at G1 phase. The different results from different lab may be due to the different cell lines we used.

How KPNA2 contributes to tumorigenesis and the development of osteosarcoma is still unclear. Yoichi Miyamoto reported that functional importin α1 (KPNA2) is located at the cell surface and may accelerate the proliferation of cancer cells by affecting FGF1 signaling in cancer cells (24). Other studies also reported that microRNAs may regulate KPNA2 expression in cancers [25]. A further study is needed to explore how KPNA2 regulates osteosarcoma in the future.

## Conclusions

In summary, our results first show that KPNA2 is overexpressed in osteosarcoma and plays critical roles in the Saos-2 osteosarcoma in vitro. This study will provide the basis for exploration of the role of KPNA2 as a therapeutic targetin osteosarcoma. Furthermore, the mechanism by which KPNA2 affects osteosarcoma cell biology requires additional exploration in the future researches.

## Acknowledgements

We thank Dr Duan Zhiqing for providing us assistance with editing the results of this study. The study was supported by the Natural Foundation of Shan Xi Province. The study was supported by the Shanxi Natural Science Fund (201603D321038;201605D211024).and The National Natural Science Foundation of China(81772867).

## List of abbreviations

ARM: armadillo
DMEM: Dulbecco’s modified Eagle’s medium
DMSO: dimethyl sulfoxide
EOC: epithelial ovarian cancer
HCS: high-content screening
IHC: immunohistochemistry
KPNA2: Karyopherin α2
NC: negative control
NPCs: nuclear pore complexes
OD: optical density
PBS: Phosphate-buffered saline
PI: propidium iodide
PVDF: polyvinylidene difluoride
qPCR: quantitative PCR
RPMI: Roswell Park Memorial Institute
RT-PCR: real-time polymerase chain reaction
SDS-PAGE: sodium dodecyl sulfate-polyacrylamide gel electrophoresis
shRNAs: short-hairpin RNAs

